# State-space Modeling Clarifies Productivity Regime Shifts of Japanese Flying Squid

**DOI:** 10.1101/2020.05.22.111088

**Authors:** Shota Nishijima, Hiroshi Kubota, Toshiki Kaga, Suguru Okamoto, Hisae Miyahara, Hiroshi Okamura

## Abstract

Regime shifts of climatic and environmental conditions potentially affect productivity of fisheries resources, posing challenging issues to stock management. The stocks of the Japanese flying squid (*Todarodes pacificus*) are suspected to suffer from regime shifts, but their detection is difficult and possibly doubtful because the nature of short-lived species readily makes the effect of regime shifts confounded with observation errors. Here we developed a new state-space assessment model to evaluate the influence of regime shifts on spawner-recruitment relationship of the Japanese flying squid. The model simultaneously estimates the population dynamics of multiple stocks that could share some life history parameters, making parameter inference stable. We demonstrate that two-time regime shifts of productivity around 1991 and 2015 caused two-to three-fold changes of maximum sustinabile yields. The model with regime shifts clarifies the relationship between fishing pressure and spawner abudance that is difficult to detect in a model with no regime shift. The state-space approach will be a promising tool to accurately assess stock status by separating recruitment process from observation errors and contribute tothe management of marine biological resources sensitive to regime shifts.

## INTRODUCTION

Ecological regime shifts cause drastic changes of ecosystem states and organisms (Scheffer et al. 2001; Kadowaki et al. 2018). In the ocean, climatic and environmental conditions drive regime shifts in productivity of fisheries stocks: average recruitment for a certain period is substantially different before and after a single year (Perälä and Kuparinen 2015; Maunder and Thorson 2019). Recent studies showed that a high proportion of stocks experienced shifts of recruitment, or nonstationary stock-recruitment relationship (Vert-Pre et al. 2013; Perälä and Kuparinen 2015; Szuwalski et al. 2015). Since fisheries production is one of the most important provisioning ecosystem services, understanding regime-shift dynamics of fisheries resources is a key issue toward the sustainable use of nature’s contribution from marine ecosystems.

Maximum sustainable yield (MSY) is an importatnt concept for the assessment and management of fish stocks around the world. International legal frameworks for sustainable fisheries, the United Nations Convension on Law of the Seas (UNCLOS) and the United Nations Fish Stocks Agreement (UNFSA), set an objective as the maintainance and restoration of populations at stock biomass that produces MSY. The Convension on Biological Diversisity (CBD) and the Sustainable Developoment Goals (SDGs) also outline the sustainable use and conservation of biological resources. Although these international circumstances require the estimation of MSY worldwide (Costello et al. 2012; Martell and Froese 2013; Punt et al. 2014; Ichinokawa et al. 2017), the existence of regime shifts makes the calculation of MSY challenging, because regime shifts are likely to generate multiple stock-recruitment relationships and thus multiple MSY-based reference points. It is suggested that although management advice should take into account recruitment variability by regime-shift-like behaviors (Vert-Pre et al. 2013; King et al. 2015), regime-based harvest control rules (HCRs) generally have high risk of overfishing when regimes are misidentified or regime shifts do not occur (A’mar et al. 2009; Szuwalski and Punt 2013). Reliable assessment on the occurrence and degree of regime shifts is needed for considering management strategies of fisheries stocks exhibiting nonstationary recruitment.

The stocks of Japanese flying squid (*Todarodes pacificus*) are considered to suffer from regime shifts in association with dynamic climatic conditions (Sakurai et al. 2002; Kidokoro et al. 2010; Kurota et al. 2020). Climatic shift in 1989 caused water temperature warm and expanded spawning areas of this species in the Sea of Japan (Sakurai et al. 2000; Kidokoro et al. 2010). Its paralarvae were likely to survive as a result of warming temperature (Sakurai et al. 2000, 2013), and thus, the catch biomass and abundance index rised dramatically since 1989 (Sakurai et al. 2002; Fig. 1). We expect, therefore, that the effect of regime shift changed spawner-recruitment relationship of this species. Furthermore, the opposite direction of shift might occur recently because the catch biomass and abundance index have been decreasing (Fig. 1; Kaga et al. 2019). Since the Japanese flying squid is preyed upon in great numbers by large fish, such as mackerels (*Scomber japonicus* and *S. australasicus*) and bullfin tuna (*Thunnus thynnus*), and marine mamamals, such as dolphins (Sakurai et al. 2013), this species has an important role in sustaining food webs in marine ecosystems. The occurrence of regime shifts in this speciesis of serious concern to fishermen and fisheries managers.

**Figure 1:**
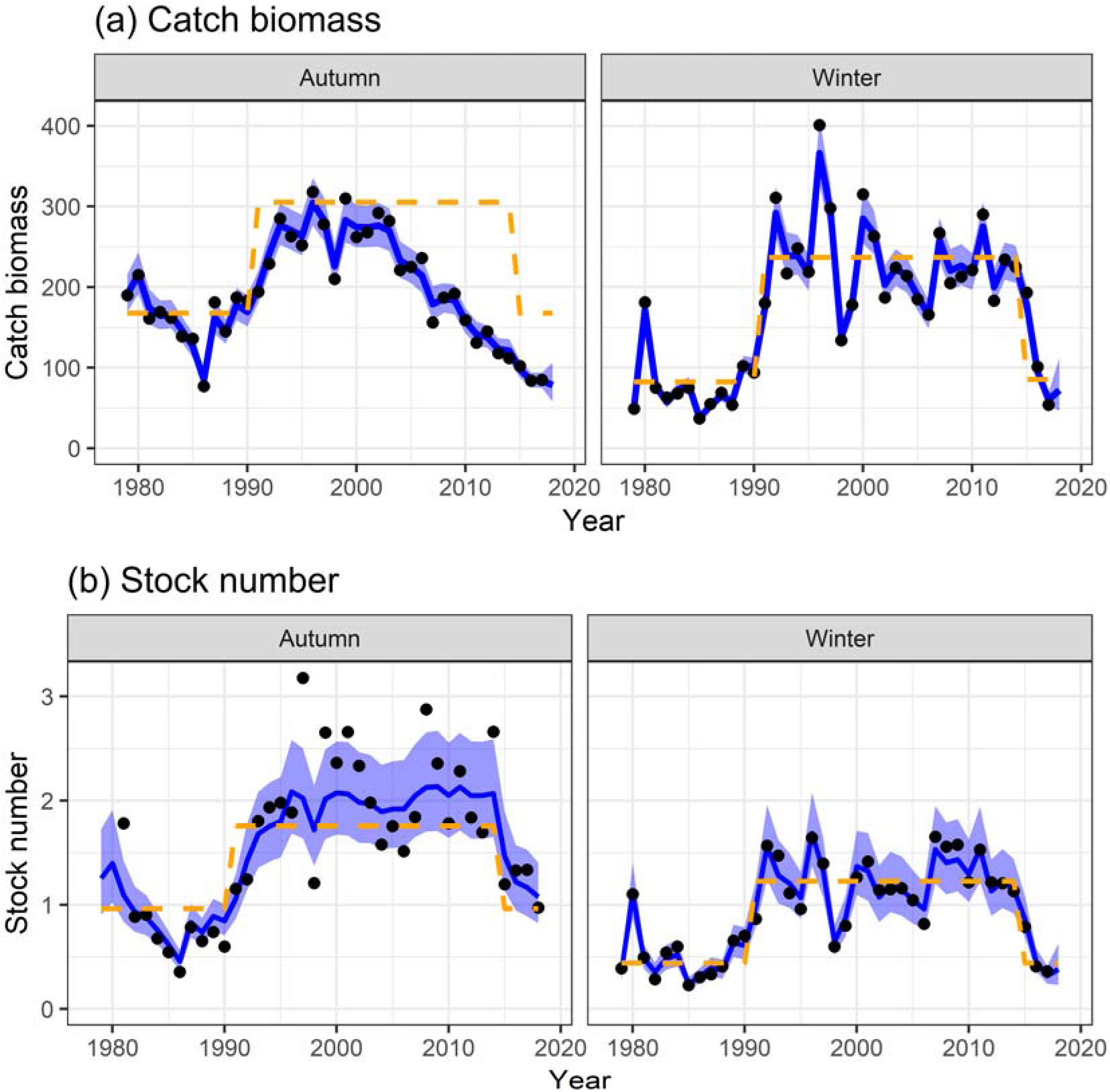
Time series of (a) catch biomass (thounsand MT) and (b) stock number (billion) for the autumn-spawning stock (left) and the winter-spawning stock (right). The black points indicate (a) observed catch biomass and (b) abundance index divided by the proportional constant (*I_i_*/*q_i_*). The blue solid lines and shadowed areas indicate point estimates and their 80% confidence intervals, respectively. The orange dashed lines indicate (a) MSY and (b) the stock number at the MSY-level equilibrium (*N_MSY_*).

Distinguishing recruitment process and observation error is important for accurately detecting regime shifts (King et al. 2015; Maunder and Thorson 2019). Recently, the state-space stock assessment models (SAM) that estimate latent variables such as abundance and fishing mortality as random effects have been developed and is effective at separately estimating process and measurement errors (Nielsen and Berg 2014; Miller and Hyun 2017; Okamura et al. 2018). However, these models are age-structured and not possible to apply to the Japanese flying squid, because its life-span is a single year. Applying population dynamics modeling to the stock assessment for a species with annual life-span is generally difficult, because one cannot track the interannual depletion process of each cohort by fishing and natural death, making parameter estimation unstable. In fact, annual stock assessment of the Japanese flying squid has been conducted based on an abundance index and not used population models (Kaga et al. 2019; Kubota et al. 2019). The stock assessment has therefore confounded measurement and process errors and been likely to fluctuate unwantedly. The calculation of MSY also increases the demand for state-space approach for the Japanese flying squid, because the estimation error in recruitment should be directly linked to the measurement error in spawner abundance (i.e., independent variable for recruitment), which could be considered appropriately by using a state-space model (Subbey et al. 2014; Brooks and Deroba 2015).

A possible solution to estimation difficulty and unstability is joint modeling of multispecies or multistocks rather than per-stock analysis (Thorson et al. 2013). Dynamics of multiple species and stocks could be partially correlated if they share environmental conditions (Thorson and Minto 2015; Thorson et al. 2016). The assumption on a species having the same life history paramter between different stocks could be valid and enable parsimonious and stably predictive modeling. Fortunately, there are two different stocks of the Japanese flying squid (autumn-spawning stock and winter-spawning stock) that have been independently assessed (Kaga et al. 2019; Kubota et al. 2019), but may have correlated dynamics (Hoshino et al. 2014).

In this article, we developed a new model for multistocks of annual life-span species, called ‘SAMUIKA’ (State-space Assessment Model Used for IKA (squid in Japanese)), to investigate whether and how regime shifts occur in productivity of the Japanse flying squid. We firstly performed intensive model selection by varying the occurrence, parameter, year, and pattern of regime shifts. We then computed MSY-based reference points from estimated spawner-recruitment relationship. Lastly, we evaluated the past stock status relative to the MSY-based reference points.

## MATERIAL AND METHODS

### Biology and fisheries of Japanese flying squid

The Japanese flying squid is one out of the nine TAC (total allowable catch) species in Japan, whose total catch are restrictly managed by output control, because it is commercially important for Japanese fisheries (5% of total Japanese catch in 2014; Kaga et al. 2017; Kubota et al. 2017). Japan has conducted annual stock assessments of autumn-spawning stock and winter-spawning stock that have diferent distributions as well as spawning seasons. The former stock is distributed in the Sea of Japan, whereas the latter is mainly distributed in the Northwest Pacific near Japan when it migrates in feeding season (Kidokoro et al. 2010). The Japanese flying squid is usually caught by jigging in Japan, but other fisheries including bottom trawling, set net, and purse seine also harvest the species especially for the winter-spawning stock (Kaga et al. 2019).

### State-space modeling

Our state-space model SAMUIKA simultaneously describes the population dynamics of autumn-spawning and winter-spawning stocks of Japanese flying squid, whose lifespan is a single year. The squid individuals that survive from natural death and fishing after recruitment become spawning adults:

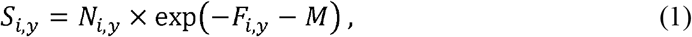

where *S_i,y_* is the number of spawning adults of stock *i* (A: autumn-spawning, W: winter-spawning) in year *y* (this definition of subscripts will be applied hereafter), *N_i,y_* is the number of recruits, or stock number. *F_i,y_* is the fishing mortality coefficient, while *M* is the natural morality coefficient and assumed to be 0.6, in accordance with the annual stock assessment (Kaga et al. 2019; Kubota et al. 2019). The natural morality corresponded to a death rate during half-year fishing season (monthly mortality coefficient was assumed to be 0.1). The interannual dynamics of fishing mortality coefficient is described by a random walk, as with age-structured state-space assessment model (Nielsen and Berg 2014):

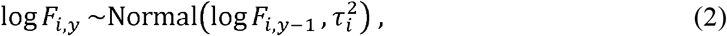

where 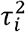 is the variance that controls the process error of random walk.

The number of recruits is expressed by the product of Beverton-Holt model and process error:

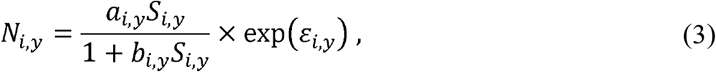

where *a_i,y_* represents the number of recruits per spawning individual when adult number approaches zero, and *b_i,y_* represents the strength of density dependence per spawning individual. We consider that the spawner-recruitment parameters *a_i,y_* and *b_i,y_* could depend not only stocks but also years because regime shifts could affect these parameters (details are shown in the next subsection). *ε_i,y_* is a deviance to the stock-recruitment curve and assumed to follow a multivariate normal distribution:

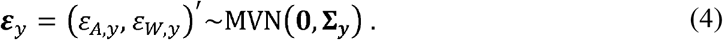

**∑**_***y***_ is a variance-covariance matrix:

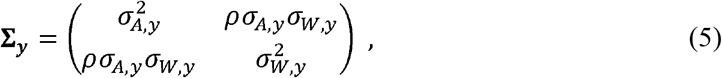

where 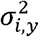 is the variance in recruitment process and *ρ* is the correlation coeffieicnt of recruitment variability between two stocks. We used the multivariate normal distribution because recruitment deviances could be correalted between autumn-spawning and winter-spawning stocks that potentially share environmental and climatic conditions. Moreover, we consider that the magnitude of recruitment variability could be different between stocks and regimes.

The following observation model was fitted to data of catch biomass and abundance index for autumn-spawing and winter-spawning stocks (Fig. 1; Kaga et al. 2019; Kubota et al. 2019). We used one time series of abundance index per stock that was used in the annual stock assessment (Kaga et al. 2019; Kubota et al. 2019). The duration of index data is 1981 to 2018 for the autumn-spawning stock and 1979 to 2017 for the winter-spawning stock (Table 1). The abundance indices were assumed to be proportional to the stock numbers with normal errors at logarithmic scale:

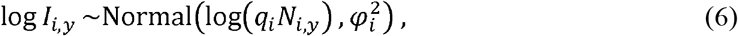

where *I_i,y_* represents an index value, *q_i_* represents a proportional constant, and 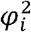 is the variance that controls the magnitude of observation error. The observed catch biomass was also followed to a normal distribution at logarithmic scale:

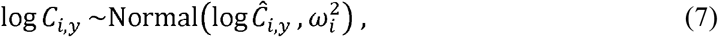

where *Ĉ_i,y_* represents a predicted catch biomass and 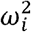 is the variance in observation error. The duration of catch data is 1979 to 2017 for both stocks (Table 1). We used the Baranov equation to obtain the predicted catch biomass:

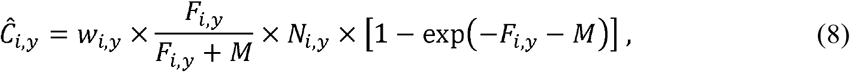

where *w_i,y_* is mean body mass per individual. We used the body weight in the annual stock assessment (Kaga et al. 2019; Kubota et al. 2019): for the autumn-spawning stock the per-capita mass is 280g during the whole period, while for the winter-spawning stock the per-capita mass is 300g before 1989 and 311g thereafter.

**Table 1:** List of symbols with their definitions, types, and constraint

| Symbol | Definition | Type<sup>1</sup> | Constraint |
| --- | --- | --- | --- |
| $I_{i,y}$ | Abundance index (1981-2018 for autumn-spawning stock, 1979-2017 for winter-spawning stock) | Data | - |
| $C_{i,y}$ | Catch biomass (1979-2017 for both stocks) | Data | - |
| $w_{i,y}$ | Per-capita body mass (1979-2017 for both stocks) | Data | - |
| $M$ | Natural mortality coefficient | Assumed | 0.6 |
| $\tau_i$ | Standard deviation of process error in random walk of fishing mortality coefficient | Fixed | Common between stocks |
| $a_{i,y}$ | Maximum number of recruits per spawning individual | Fixed | At least either $a_{i,y}$ or $b_{i,y}$ is common between stocks and regimes |
| $b_{i,y}$ | Strength of density dependence per spawning individual | Fixed | |
| $\sigma_{i,y}$ | Standard deviation of recruitment variability | Fixed | - |
| $\rho$ | Correlation coefficient of recruitment variability between stocks | Fixed | - |
| $q_{i,j}$ | Proportional constant for abundance index | Fixed | - |
| $\varphi_i$ | Standard deviation of observation error for abundance index | Fixed | - |
| $\omega_i$ | Standard deviation of observation error for catch biomass | Fixed | Common between stocks |
| $F_{i,y}$ | Fishing mortality coefficient | Random | - |
| $N_{i,y}$ | Number of recruits (stock number) | Random | - |
| $S_{i,y}$ | Number of spawning adults | Derived | - |
| $\varepsilon_{i,y}$ | Recruitment deviation to stock-recruitment relationship | Derived | - |
| $i$ | Stock (A: autumn-spawning, W: winter-spawning) | Subscript | - |
| y | Fishing year | Subscript | - |

1: "Fixed", "Random", and "Derived" are the types of parameters.

### Parameter estimation and model selection

We estimated the parameters of fixed and random effects (Table 1) using the maximum likelihood method via template model builder (TMB, Kristensen et al. 2016). TMB enables the estimation of many random effects using the Laplace approximation and automatic differentiation (Kristensen et al. 2016). Because the random effects were estimated at the logarithmic scale, we applied a generic method for bias correction for the mean of random effects (Thorson and Kristensen 2016). The source code and data are made available as an R package at GitHub (https://github.com/ShotaNishijima/messir).

TMB enables fast optimization of hierarchical models including complex random effects (Kristensen et al. 2016). By utilizing this advantage, we analyzed a number of models having different assumptions on recruitment and performed model selection based on AICc (Burnham and Anderson 2002). We found that assuming that both *a_i,y_* and *b_i,y_* were independent between stocks caused false convergence or unrealistic, extremely-large abundance estimates, suggesting that estimation is unfeasible only from one-stock information. We therefore assumed that at least either *a_i,y_* or *b_i,y_* must be a common value between stocks. We also assumed that a regime shift occurred simultaneously for both stocks and changed either *a_i,y_* or *b_i,y_* in the spawner-recruitment relationship. This assumption was made because a previous study suggeted the shift in climatic conditions changed spawining areas and stock abundances for both autumn-spawning and winter-spawning stocks (Sakurai et al. 2000). The changes in *a_i,y_* and *b_i,y_* both caused the change in productivity, and therefore, were likely to be confounded (Maunder and Thorson 2019). The variation in *a_i,y_* changes both maximum recruits per spawner and maximum recruitment, and thus affects expected recruitment at both high and low spawning abundances. On the other hand, the variation in *b_i,y_* changes maximum recruitment, but not maximum recruits per spawner, and thus affects expected recruitments at high spawning abundance.

To reduce the number of analyzed models, we assumed that the parameter that are independent of stocks could change in response to a regime shift by considering that the spawner-recruitment parameter that is different between stock is also likely to be differnt among regimes; when the parameter *a_i,y_* (or *b_i,y_*) was different between stocks, *a_i,y_* (or *b_i,y_*) could be different among regimes. When the same parameter values *a_i,y_* and *b_i,y_* were shared between stocks, we assumed that either *a_i,y_* or *b_i,y_* changed due to a regime shift. We thus made seven types of assumptions: (1) both paramters *a_i,y_* and *b_i,y_* were common between stocks and no regime shift occured; (2) both paramters *a_i,y_* and *b_i,y_* were common between stocks and a regime shift changed the parameter *a_i,y_*; (3) both paramters *a_i,y_* and *b_i,y_* were common between stocks and a regime shift changed the parameter *b_i,y_*; (4) the parameter *a_i,y_* were different between stocks (*b_i,y_* were common) and no regime shift occured; (5) the parameter *b_i,y_* were different between stocks (*a_i,y_* were common) and no regime shift occured; (6) the parameter *a_i,y_* were different between stocks (*b_i,y_* were common) and a regime shift changed the parameter *a_i,y_*; and (7) the parameter *b_i,y_* were different between stocks (*a_i,y_* were common) and a regime shift changed the parameter *b_,y_*.

Previous studies suggests that a regime shift from low to high state occurred around 1989 (Sakurai et al. 2002; Kidokoro et al. 2010), and another regime shift possibly occurs in recent years (Kaga et al. 2019). We considered three pattens of regime shifts: (i) a regime shift occurred once in a year between 1987-1991 (A→B); (ii) regime shifts occurred twice in a year between 1987-1991 and in a year between 2013-2017, respctively, and the second regime shift reverted the first state (A→B→A); and (iii) regime shifts occurred twice in a year between 1987-1991 and in a year between 2013-2017, respctively, and the second regime shift brought a third state (A >B >C). The first patten had five cases having different shifting years, and the second and third patterns had 25 cases (= 5×5), and thus we analyzed 55 cases (5+15+25) in the models with regime shift(s). We had four out of the seven types that assumed regime shift(s) in the previous paragraph (2, 3, 6 and 7). The other three types with no regime shift had only one case respectively (1,4, and 5). We analyzed 223 models (= 55×4+3) with different assumptions in total.

We futher assumed the parameter 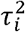 (variance representing process error of fishing mortality coefficient) and 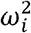 (variance representing observation error of catch) were common between stocks (Table 1). This is because a preliminary analysis showed that estimated values of these parameters varied little between stocks and assuming a common value between stocks had lower AICc (Burnham and Anderson 2002) than assuming different values in a preliminary analysis. We excluded 68 models that caused estimation error or convergence problem from results; remaining 155 (223–68) models achieved successful estimation and convergence. We calculated AICc of each of successfully-converged models from maximum likelihood, sample size, and the number of fixed parameters shown in Table 1.

### MSY-based reference points

We calculated derived parameters and biological reference points from estimated stock-recruitment relaltionships (Eq. 3). First, we obtained the steepness *h*, or the fraction of recruitment from unfished population obtained when the spawning stock is 20% of its unfished level (Mangel et al. 2010):

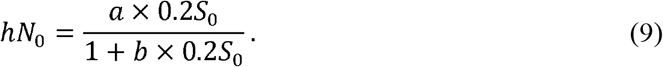

*N*_0_ and *S*_0_ are the unfished numbers of recruits and spawners, respectively, which can be obtained from the intersection of the spawner-recruitment relationship and the replacement line (*y* = exp(*M*)×*x* in this case):

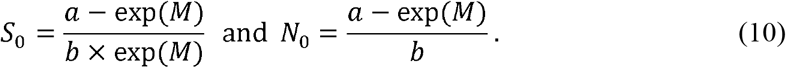

*h* = 1 means that the recruitment is completely driven by environments, whereas *h* = 0.2 means the proportional relationship between spawners and recruits. The steepness, therefore, represents the resilience of a species to harvesting: a high steepness indicated high resilience, and vice versa (Mangel et al. 2010). In this study, we can calculate the steepness by substituting the equation (10) into the equation (9):

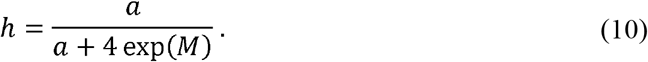

It is worth noting that the steepness depends on the parameter *a* but not *b*.

We then calculataed MSY-based reference points. The amount of surplus production reaches at maximum when the difference between spawner-recruitment relationship and replacement line is the largest:

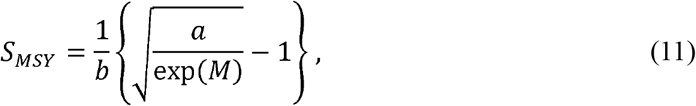

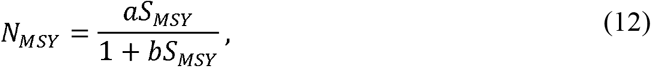

and *F_MSY_* = log(*N_MSY_*)−log(*S_MSY_*)−*M*. MSY was calculated by substituting *N_MSY_* and *F_MSY_* into Eq. 8. We compared the estimates with these MSY-based reference points to evaluate the stock status in the past. The steepness and MSY-based reference points were computed for each paramter set of spawner-recruitment relationship when models with different parameters between stocks and/or regimes were selected.

## RESULTS

### Model selection

Model selection showed that top models with lower AICc had two-time regime shifts around 1991 and 2015 (Table 2). The models with one-time regime shift had ΔAICc of 21.7 or larger. The no-regime models had ΔAICc of 34.1 or larger, and were ranked as the worst among the 155 models having successful convergence. For comparison, the results of a no-regime model are shown in Supporting Information. The best five models assumed different parameter values of the strength of density dependence (*b_i,y_*) between stocks and regimes, rather than the maximum number of recruits per spawning individual (*a_i,y_*). These five had different years of regime shifts, but the best model, which assumed 1991 and 2015 as shifting years, had significantly lower ΔAICc than the other models (ΔAICc of 4.2 or larger). The best three models assumed that the second regime shift went back to the first regime (A→B→A), but the forth model with ΔAICc of 5.8 assumed different regimes between the first and third ones (A→B→C). The sixth model with ΔAICc of 5.9 had different parameter values of the maximum number of recruits per spawning individual (*a_i,y_*), but a common value for the strength of density dependence (*b_i,y_*).

**Table 2:**
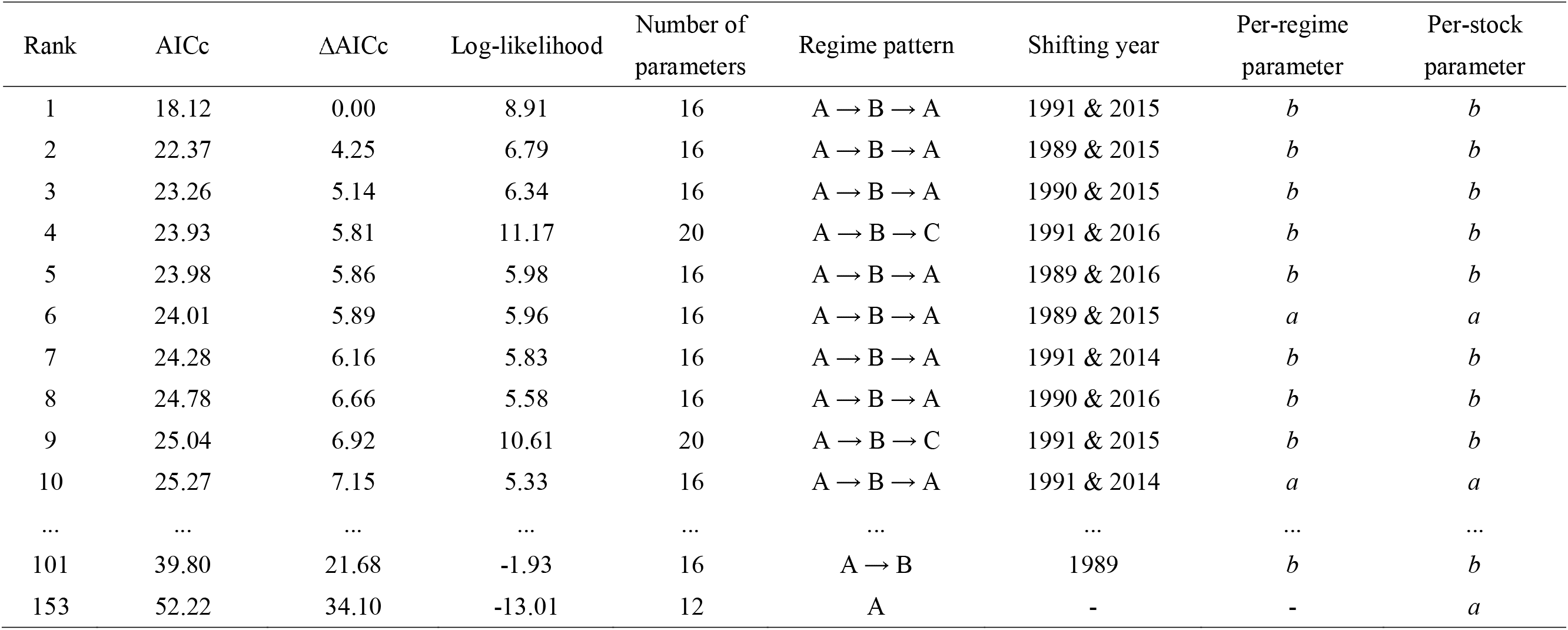
Model selection table for top ten models, the best model with one-time regime-shift, and the best model with no regime

### Fit to observation

The estimated temporal patterns of stock number were smoother than the temporal dynamics of abundance indices especially for the autumn-spawning stock (Fig. 1b). The stock number was kept at the low level during the first decade, and then abruptly increased since the 1990s. This ‘high’ regime continued to a middle of the 2010s and thereafter, the ‘low’ regime came back. The estaimtes of catch biomass were well fitted to its observed values (Fig. 1a).

### Recruitment productivity

The spawner-recruitment relationships were clearly distinct between the regimes in the best model (Fig. 2). The Japanese flying squid belonged to the low regime in the 1980s, and thereafter, moved to the high regime. The productivity then decreased to the low regime in 2015. The magnitude of regime shift was largher for the winter-spawning stock (Fig. 2b) than for the autumn-spawning stock (Fig. 2a). For the autumn-spawning stock, the MSY in the high regime (305 thousand MT) was 1.8 times larger than that in the low regime (167 thousand MT); for the winter-spawning stock, by contrast, the MSY in the high regime (237 thousand MT) was 2.8 times larger than that in the low regime (85 thousand MT) (Fig. 1a).

**Figure 2:**
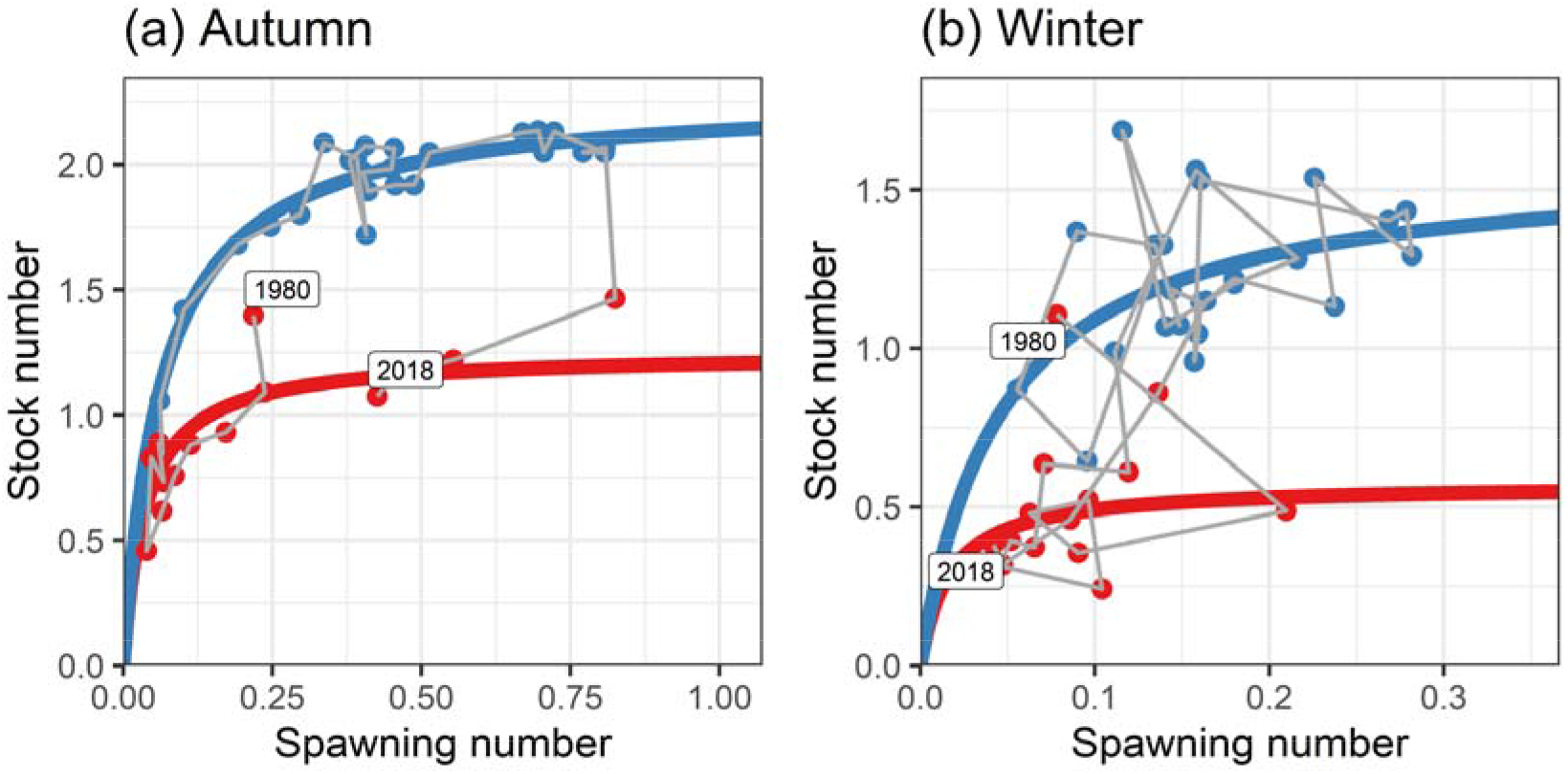
Spawner-recruitment relationships of (a) the autumn-spawning stock and (b) the winter-spawning stock. The red and blue lines indicates the low and high regimes, respectively.

The spawner-recruitment relationship with regime shifts was considerably different from that with no regime shift. The spawner-recruitment relationship became close to a proportional relationship when we ignored regime shifts (Fig. S2 in Supporting Information). The steepness was 0.83 in the best model with regime shifts, but 0.35 in the model with no regime. This indicates that incorporating regime shifts made the Japanese flying squid more resilient to harvesting.

Recruitment variability was higher in the winter-spawning stock than in the autumn-spawning stock (Fig. 2). The recruitment variablity was moderately correlated between the stocks (*ρ* = 0.69).

### Fishing impact on spawners

The temporal patterns of fishing mortality greatly difffered between stocks. For the autumn-spawning stock, the fishing mortality coefficient was higher than *F_MSY_* in the 1980s, but gradually decreased to a much lower level than *F_MSY_* (Fig. 3a). As a result, the spawning number of autumn-spawning stock was kept at a higher level than *S_MSY_* (Fig. 3b). The relationship between the relative fishing mortality coefficient (*F*/*F_MSY_*) and the relative spawning abundance (*S*/*S_MSY_*) showed a clear negative association (Fig. 4).

**Figure 3:**
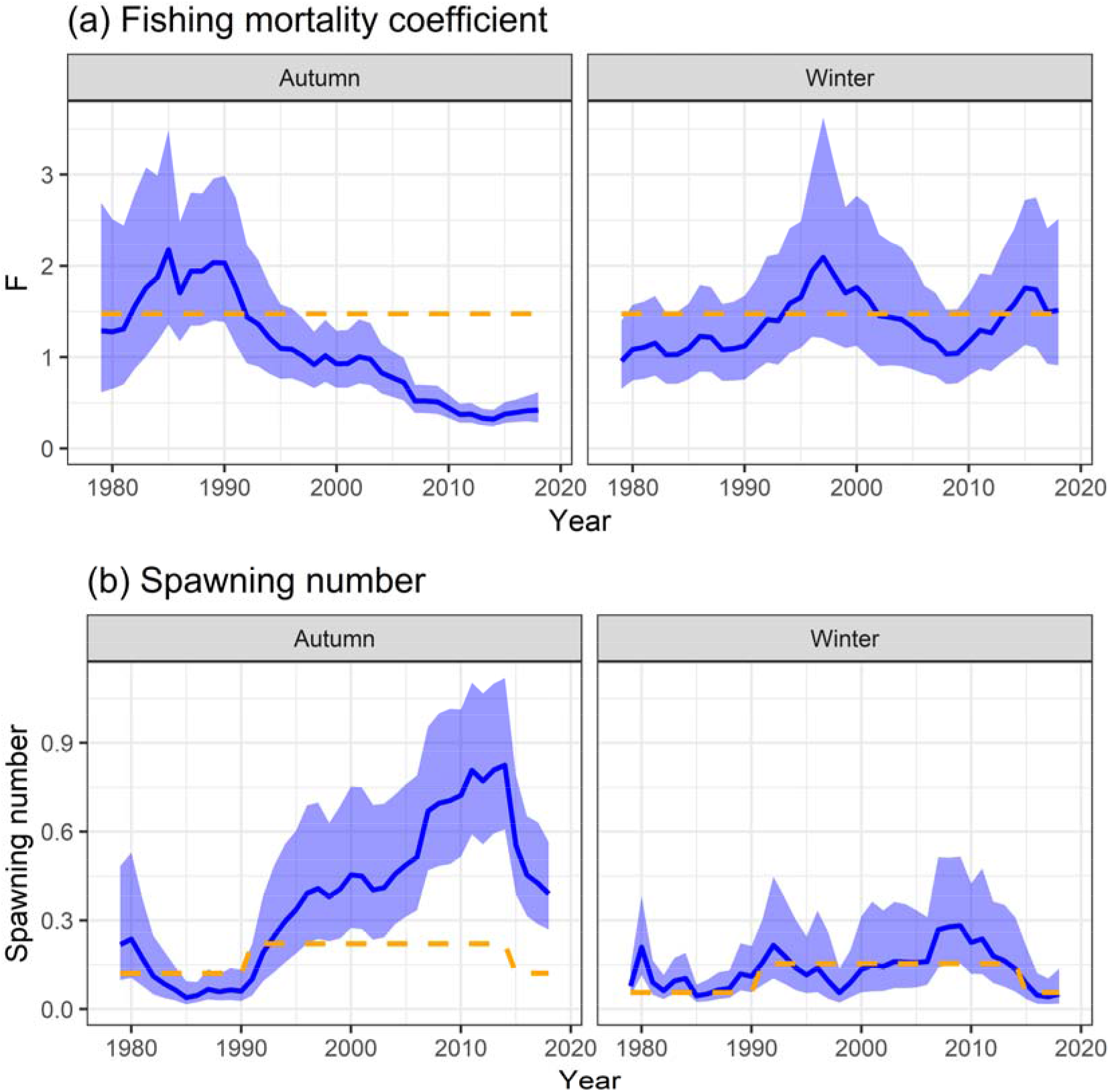
Time series of (a) fishing mortality coefficient and and (b) spawning number (billion) for the autumn-spawning stock (left) and the winter-spawning stock (right). The blue solid lines and shadowed areas indicate point estimates and their 80% confidence intervals, respectively. The orange dashed lines indicate the MSY-level equilibrium (*F_MSY_* and *S_MSY_*).

**Figure 4:**
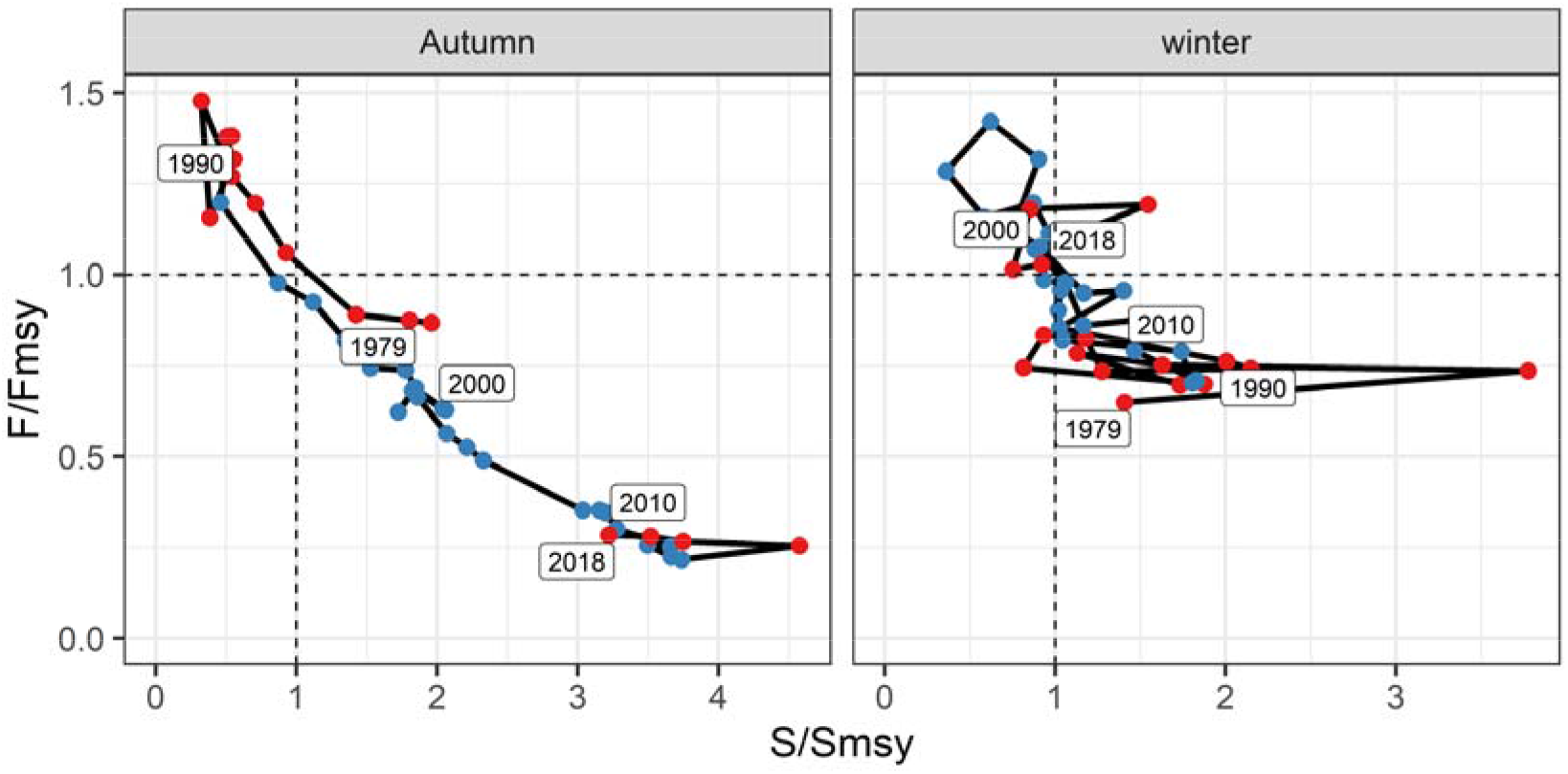
Relationships between fishing mortality coefficient and spawner abundance relative to the MSY-based reference points for the autumn-spawning stock (left) and the winter-spawning stock (right). The red and blue lines indicates the low- and high-regimes, respectively.

For the winter-spawning stock, the fishing mortality coefficient wandered around the level of *F_MSY_* (Fig. 3a). The fishing pressure was lower than *F_MSY_* up to 1993, but became higher from 1994 to 2001. Thereafter, although the fishing impact was kept at a lower level than *F_MSY_* up to 2013, it became higher again in recent years. The spawning number thus stayed around the MSY level (*S_MSY_*) (Fig. 3b). The relative spawning abundance (*S*/*S_MSY_*) exhibited an opposite trend of the relative fishing mortality coefficient (*F*/*F_MSY_*) (Fig. 4).

It is noteworthy that *F_MSY_* was constant between regimes and between stocks (Fig. 3a). This is because model selection favored the best model that shared the maximum number of rectuits per individuals (*a_i,y_*), rather than the strength of density dependence (*b_i,y_*), and *F_MSY_* depended only on *a_i,y_*.

The patterns of fishing pressure and spawner abundance were substantially different depending on whether we considered regime shifts or not, especially for the winter-spawning stock. If we had ignored regime shifts, the fishing mortality would have exceeded *F_MSY_* (i.e., overfishing) and the spawning abundance would have been lower than *S_MSY_* (overfished) for the whole period of the winter-spawning stock (Fig. S3 in Supporting Information). The relationship between the relative fishing mortality coefficient (*F*/*F_MSY_*) and the relative spawning abundance (*S*/*S_MSY_*) was unclear in the model without regime (Fig. S4 in Supporting Information).

## DISCUSSION

The state-space assessment model clarified that productivity regime shifts of the Japanese flying squid occurred twice during the analyzed period. Previous studies based on field surveys showed that the climatic shift from cool to warm condition caused the expansion of spawning areas of this species around 1989 (Sakurai et al. 2000, 2013; Kidokoro et al. 2010). The current study provides another line of supportive evidence of regime shift for the Japanese flying squid, using population dynamics modeling. Although the climatic shift was recognized to occur in 1989 (Yasunaka and Hanawa 2002), our results showed that the top model had the shifting year of 1991 (Table 2). This might suggest a time lag of biological response of productivity to the climatic effect or be a statistical artifact because the second top model selected 1989 as the shifting year. The top models suggested that the second regime shift occurred around 2015 and the current state was identical to that in the 1980s, longer time-series data would be needed to decide whether the current regime is truly the same as the 1980s. Top models favored different parameter values of the strength of density dependence, rather than maximum recruits per capita spawner, between regimes (Table 2). Combined with biological studies showing that the survival rate of paralarvae varied in response to climatic regimes (Sakurai et al. 2000, 2013), climate-driven regime shifts may affect a density-dependent survival of paralarvae. Because TMB enables much faster parameter inference than Bayesian MCMC algorithm, it is now easier than ever to analyze a number of hierarchical models and perform intensive model selection like this study. Random-effect models will be increasingly applied to various kinds of stock assessment modeling (Thorson and Minto 2015).

SAMUIKA is a novel state-space stock assessment model in terms of multistock modeling of annual life-span species. To check estimability, we conducted a simple simulation test as an additional analysis by generating bootstrap data from estimated models (Supporting Information). Results showed that the best model with the lowest AICc could obtain almost unbiased estimates in abundances and fishing mortalities (Fig. S5 in Supporting Information). However, the no-regime model obtained seriously-biased estimates: overestimation of abundances and underestimation of fishing mortalities (Fig. S6 in Supporting Information). This bias was caused because the no-regime model had larger estimation uncertainty than the regime-shift model and ignoring regime shifts was likely to mask fishing impacts (see Supporting Information for details). Indeed, the regime-shift model estimated lower abundances and higher fishing mortality coefficients than the no-regime model, suggesting that the regime-shift model possibly obtained estimates closer to true values. Incorporating productivity regime shifts into the assessment of the Japanese flying squid fundamentally important for accurately estimating stock status.

Our concern on the model results is that estimation uncertainty in spawner abundance was larger than that in recruitment (stock) abundance: average coefficient of variation was 0.18 for stock numbers but 0.55 for spawning numbers. This is because the used abundance indices were of recruitment abundance, but not spawner abundance. The large uncertainty in spawner abundance may be problematic, because spawner abundance is usually employed to judge stock status and its uncertainty is directly linked to the reliability of stock assessment. Developing an abundance index for spawners is an important future task toward more robust estimation.

Regime shifts caused twofold and threefold changes of MSY to the autumn-spawning stock and the winter-spawning stock, respectively (Fig. 1). A reason for the larger difference of MSY of the winter-spawning stock is that it migrates in large areas off east coast of Japan in the Northwest Pacific including high seas, where the Kuroshio and Oyashio Currents cause enormous decadal variation of environmental factors (Yatsu et al. 2013). Compared to small pelagic fishes, however, the magnitude of regime shifts is smaller for the stocks of the Japanese flying squid; for example, the Pacific stock of the Japanese sardine (*Sardinops melanostictus*) has a 13-fold difference of MSY between regimes (S. Furuichi, personal comminucation). We recognize, therefore, that the Japanese flying squid probably exhibits the regime shift of productivity, but its intensity is not so large.

Two major differences between the results with and without regime shifts can be seen in the relationship between fishing mortality coefficient and spawner abundance relative to the MSY-based reference points (Fig. 4 vs. Fig. S4 in Supporting Information). First, the model with regime shifts showed a clearer negative correlation between the relative fishing mortality. This suggests that ignoring regime shifts is likely to obscure the impact of fishing and incorporating environmentally-driven productivity shifts can greatly change our view of fishing influences on stock status. Second, the stock status is more likely to be overfishing and overfished in the model with no regime shift than in the model with regime shifts. This is because the no-regime model presumed that the variation of recruitment was caused by the variation of spawner abundance, rather than regime shifts, causing lower steepness and resilience to fishing (Fig. 2 vs. Fig. S2 in Supporting Information). Accordingly, one may wrongly declare overfishing and/or overfished of a stock, if one ignores truly-occurring regime shifts. A similar result is obtained from a previous simulation study testing the effectiveness of regime-based HCRs (Szuwalski and Punt 2013).

Although our model has demonstrated the occurrence of regime shifts for the squid stocks, whether we should choose a regime-based HCR still remains uncertain. Previous studies presented two risky situations of overfishing (Szuwalski and Punt 2013; King et al. 2015): (1) one wrongly applies a HCR for high regime when one overlooks shifting from high to low regime; and (2) one wrongly applies a regime-bas ed HCR when regime shifts do not occur actually. These risks may not be high for the stocks of Japanese flying squid, however. Our estimated *F_MSY_* are common between the high and low regimes (Fig. 3), and setting the *F_MSY_* as a limit reference point will possibly avoid overfishing even if one overlooks a high-to-low regime shift. In addition, the estimates of stock abundance in the regime-shift model were smaller than those in the no-regime model and, therefore, the estimated MSY in the high regime was not larger than that in the no-regime model for the winter-spawning stock, although this was not the case for the autumn-spawning stock (Fig. 1 vs. Fig. S1 in Supporting Information). The risk of overfishing may thus be sufficiently low even if applying regime-based HCRs to this species. On the other hand, we also expect that there is no sufficient profit (i.e., increased catch) of using regime-based HCRs, because the magnitude of regime shifts is not large. Management strategy evaluation (MSE) will be useful to make a judgement on this indecisive debate. The state-space model can assist the MSE as not only assessment model but also operating model.

Our state-space modeling highlights a future direction of fisheries stock assessments. Currently, stock-recruitment relationships have sometimes been estimated by ex-post analyses using the estimates in stock assessment as fixed like observed data to detect nonstationary dynamics (Vert-Pre et al. 2013; Szuwalski et al. 2015; Kurota et al. 2020). However, abundance estimates could vary depending on whether regime shifts are incoporated into assessment models, as demonstrated by this study, and it is ideal to estiamte a stock-recruitment relationship within stock assessment models (Subbey et al. 2014; Brooks and Deroba 2015). The state-space approach can be extended by two ways. First is to incoroporate environmental effects on recruitment productivity, which can be alternative to regime-shift models and has a potential to improve the ability of future projection of stock dynamics (King et al. 2015; Maunder and Thorson 2019). Second, joint modeling of multispecies will be feasible by inferring interspecific correlation in population dynamics. Although a multispecies spatio-temporal model has recently been developed (Thorson 2019), multispecies state-space assessment models are still rare (Thorson and Minto 2015), but will be informative for evaluating and mechanistically understanding fish comminities. These two ways correspond to ecosystem- and community-based approaches. Such approaches will play an important role in evaluating provisioning ecosystem services from fisheries production as a whole, because single-species assessment is inefficient and may be insufficient for whole-scale evaluation of ecosystem services. Integrating multispecies and environmental effects into state-space assessment models will contirubte to the understanding of community dynamics and the sustainable use of marine ecosystem services.

## ACKNOWLEDGEMENTS

This study was funded by JSPS KAKENHI Grant Number 19K15905. This study was partially supported by grants for marine fisheries stock assessment and evaluation in Japanese waters from the Fisheries Agency and the Fisheries Research and Education Agency of Japan.

