## Supporting Information for "State-space Modeling Clarifies Productivity Regime Shifts of Japanese Flying Squid"

Supporting Information for “**State-space Modeling Clarifies Productivity Regime Shifts of Japanese Flying Squid**” by Nishijima et al.

### **Appendix A: Results of no-regime model**

Here we show the results of a model with no regime shift for comparison. Among the models with no regime shift, the models that had different values of the parameter *a* (ΔAICc = 34.10) or *b* (ΔAICc = 36.97) between stocks had lower AICc than the model had identical values of the parameters *a* and *b* between stocks (ΔAICc = 39.40). However, the models with different values of *a* or *b* between stocks estimated extremely low fishing mortality coefficients (F ≈ 0) and extremely high abundances. We considered the estimates were unrealistic and, therefore, showed the results with identical parameter values of *a* and *b* as the no-regime model. Figs. S1-4 below correspond to Figs 1-4 in the main text for the best model with regime shifts.

There were five major differences between the regime-shift model and the no-regime model. First, abundance estimates were higher in the no-regime model than the regime-shift model and the estimates of fishing mortality coefficients were lower in the no-regime model (Figs. S1, S3). Second, the coefficients of variation of estimated abundance and fishing mortality coefficients were larger in the no-regime model than in the regime-shift model (Figs. S1, S3), suggesting that the no-regime model had larger estimation uncertainty. Third, the spawner-recruit relationship was more linear and had lower steepness in the no-regime model than in the regime-shift model (Fig. S2), suggesting lower resilience to harvesting in the no-regime model. Forth, the stock status was more likely to be overfishing or overfished in the no-regime model than in the regime-shift model (Fig. S4). Lastly, the association between fishing impacts and spawner abundances was unclearer in the no-regime model than in the regime-shift model (Fig. S4). Whether one incorporates regime shifts had a large influence on the stock assessment of the Japanese flying squid.

Figure S1:

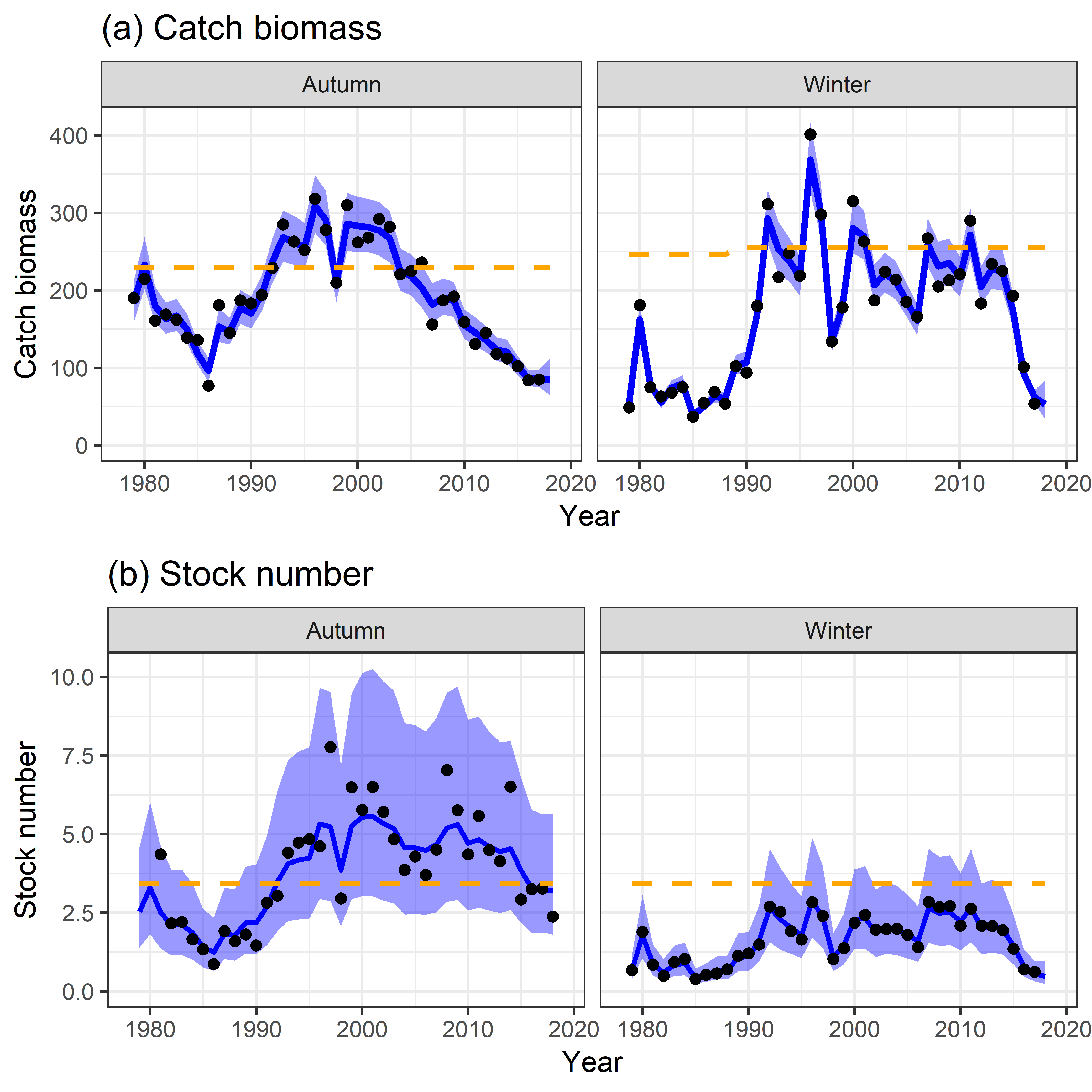

Time series of (a) catch biomass (thounsand meric ton) and (b) stock number (billion) for the autumn-spawning stock (left) and the winter-spawning stock (right). The black points indicate (a) observed catch biomass and (b) abundance index divided by the proportional constant (*I_i_*/*q_i_*). The blue solid lines and shadowed areas indicate point estimates and their 80% confidence intervals, respectively, obtained by the no-regime model. The orange dashed lines indicate (a) MSY and (b) the stock number at the MSY-level equilibrium (*N_MSY_*).

*Figure S2:*

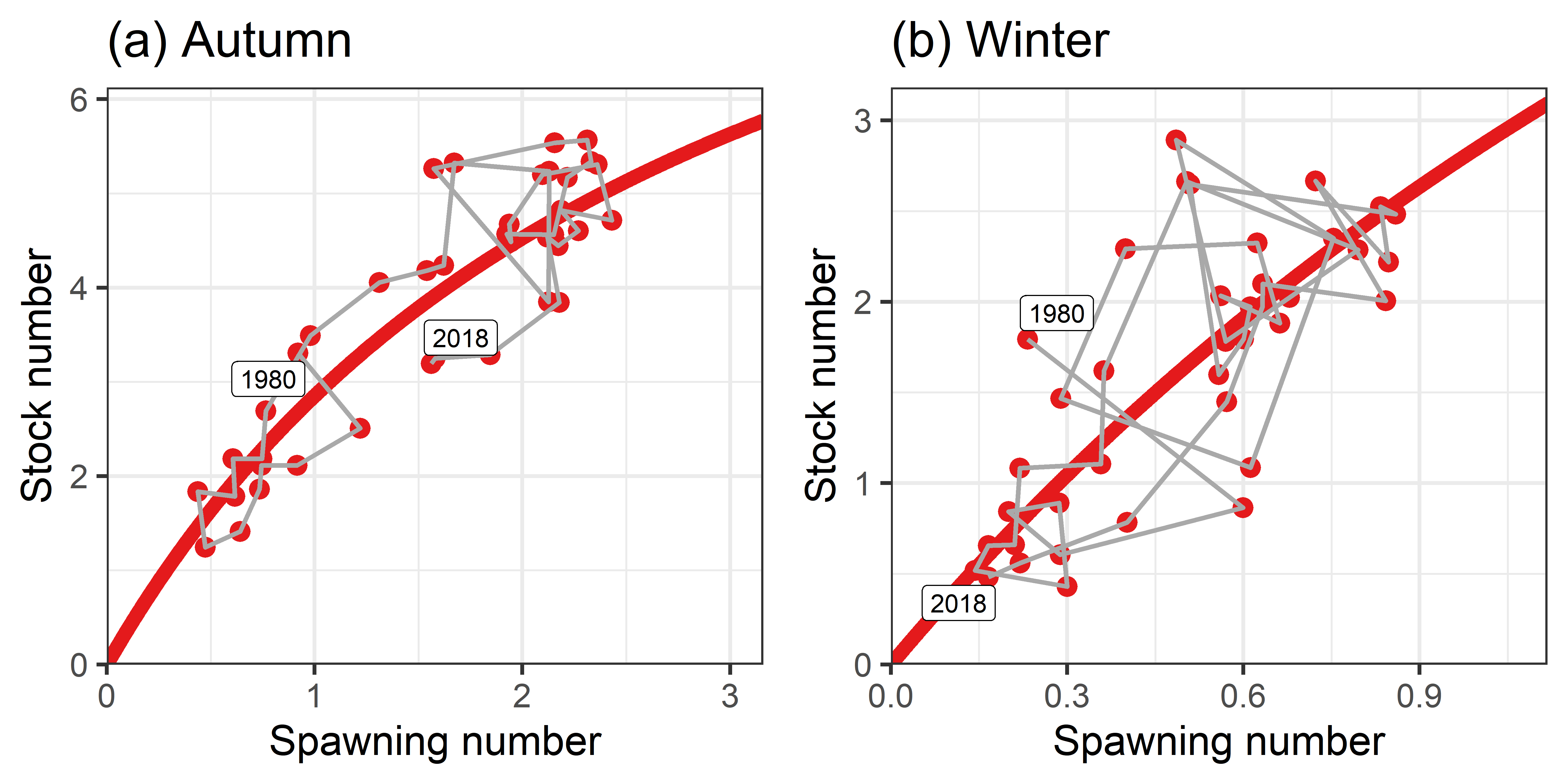

Spawner-recruitment relationships of (a) the autumn-spawning stock and (b) the winter-spawning stock, estmiated by the no-regime model.

*Figure S3:*

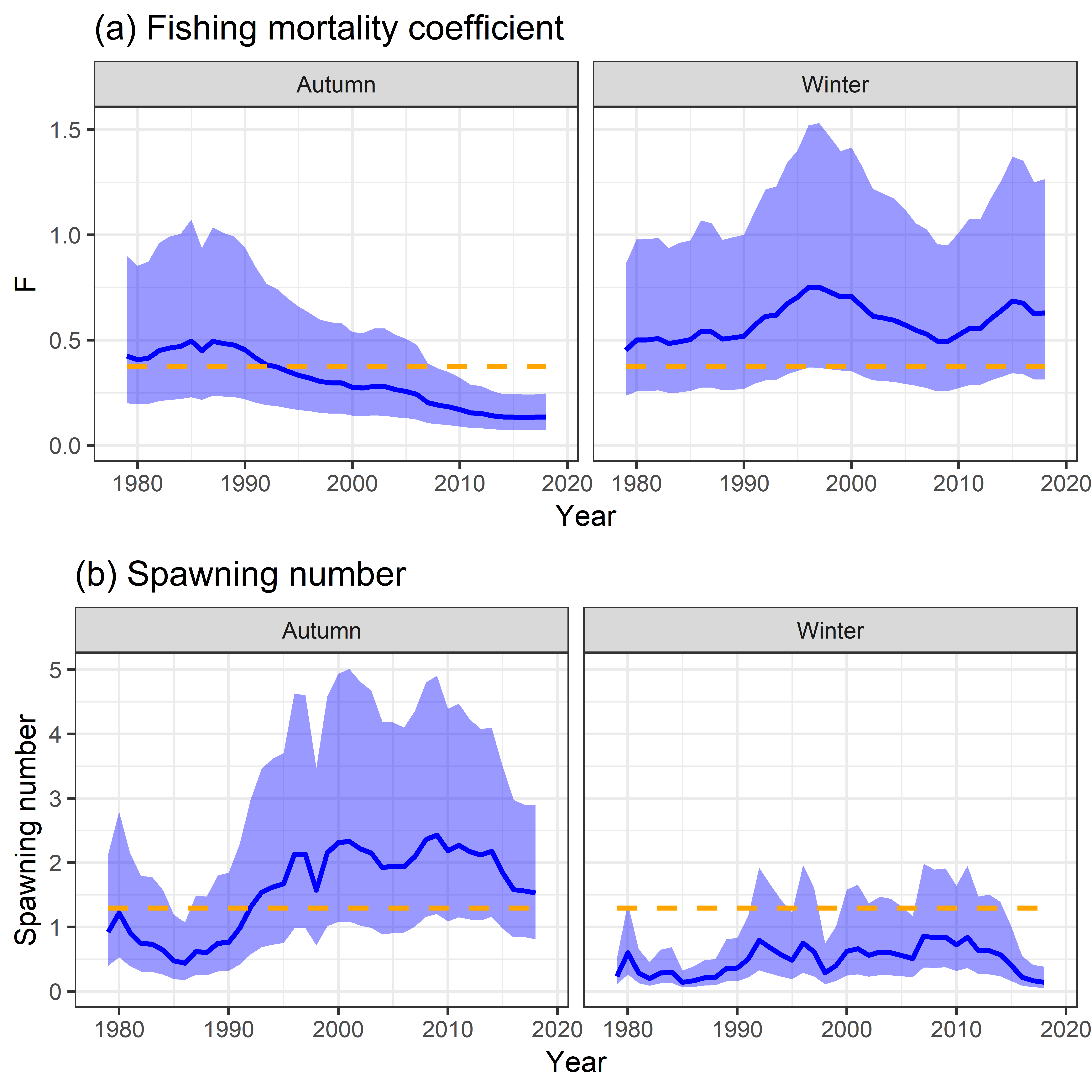

Time series of (a) fishing mortality coefficient and and (b) spawning number (billion) for the autumn-spawning stock (left) and the winter-spawning stock (right). The blue solid lines and shadowed areas indicate point estimates and their 80% confidence intervals, respectively, obtained by the no-regime model. The orange dashed lines indicate the MSY-level equilibrium (*F_MSY_* and *S_MSY_*).

*Figure S4:*

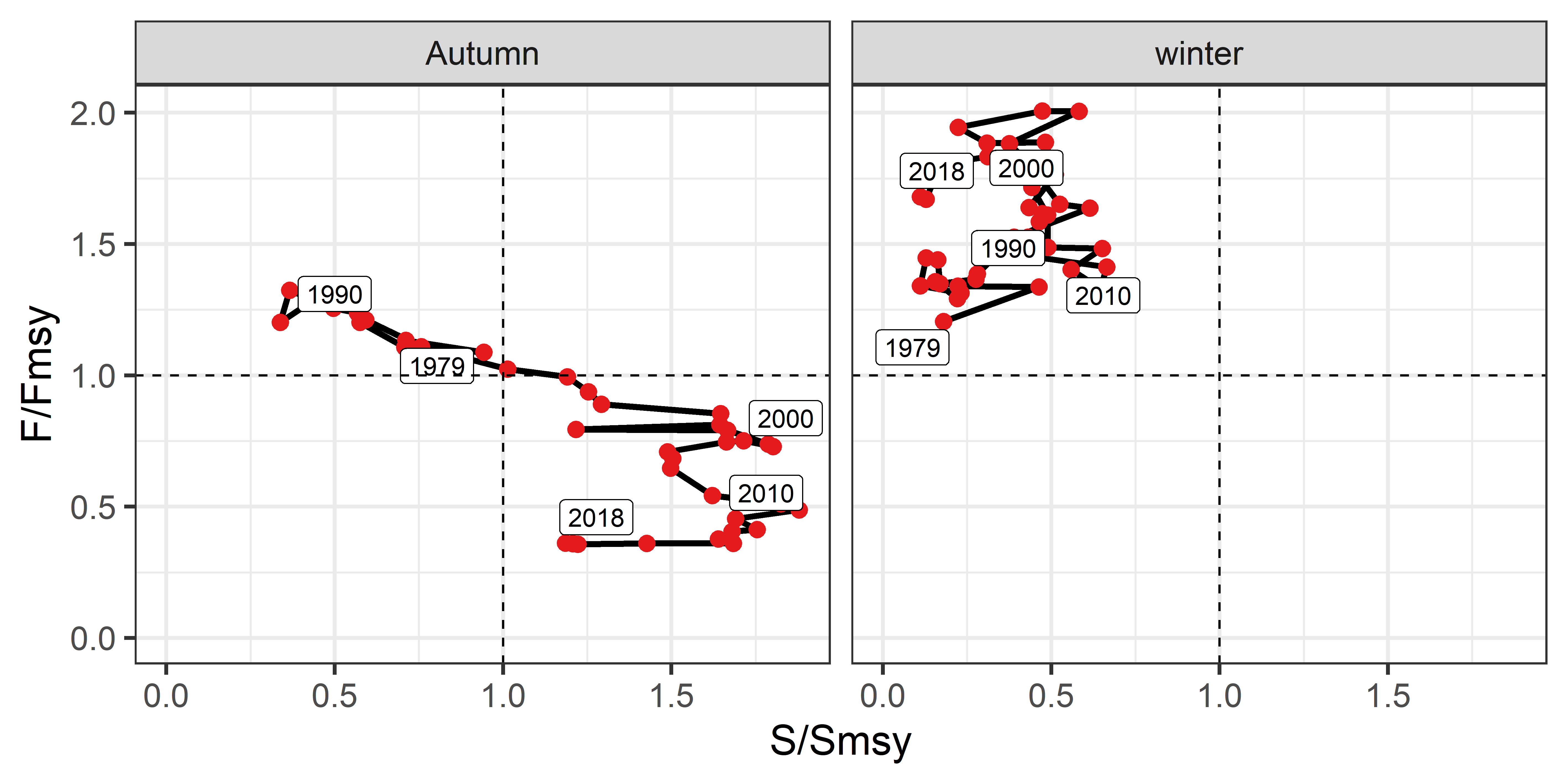

Relationships between fishing mortality coefficient and spawner abundance relative to the MSY-based reference points for the autumn-spawning stock (left) and the winter-spawning stock (right), estimated by the no-regime model

### **Appendix B: Simulation testing of estimation bias**

We conducted a simple simulation test to evaluate estimability of the state-space assessment model. We generated bootstrapped data of abundance indices and catch biomass from the equations (6) and (7) by assuming that estimated models were true:

$$\log I_{i,y}^{B}\sim\mathrm{Normal}\left( \log\left( \hat{q}_{i}\hat{N}_{i,y} \right), \hat{\varphi}_{i}^{2} \right) ,$$

(S1)

$$\log C_{i,y}^{B}\sim\mathrm{Normal}\left( \log\hat{C}_{i,y}, \hat{\omega}_{i}^{2} \right) ,$$

(S2)

where the superscript *B* represents bootstrapped data and the hat represents the estimated value. We used the best model with the lowest AICc (the regime-shift model) and no-regime model shown in Appendix A. We estimated parameters from the bootstrapped data generated from these models and replicated this trial 200 times per model.

Bootstrap results were greatly different between the regime-shift model and no-regime model. The regime-shift model obtained almos unbiased estimates of stock numbers and fishing mortality coefficients with small confidence intervals, although bootstrap estimates of fishing mortality coefficients for the autumn-spawning stocks were slightly lower than the point estimates (Fig. S5). However, the no-regime model obtained seriously biased estimates: bootstrap estimates of stock numbers were much higher than the point estimates, while bootstrap estimates of fishing mortality coefficients were much lower than the point estimates (Fig. S5). This result suggests that the point estimates of no-regime model were overestimation for stock numbers and underestimation for fishing mortality coefficients, because bootstrap estimates had these directional biases against true values (point estimates). This bias possibly derived from the instability of parameter estimation in the no-regime model; the proportional constant, *q_i_*, which a scaling factor determining the levels of abundance and fishing pressure, had much larger variance and bias in the no-regime model than in the regime-shift model (Fig. S7). Ignoring regime shifts obscured the impact of fishing on spawners (Fig. S4), which was likely to cause the estimation instability and the underestimation of fishing mortality coefficients and thus the overestimatino of stock numbers. A condition for this model to have sufficient estimability is threfore that fishing pressure must be high to a certain degree. As the point estimates of stock numbers and fishing mortality coeffieicnts were lower and higher, respectively, in the regime-shift model than in the no-regime model, it is suggested that the regime-shift model could more unbiasedly estimate stock status of the Japanese flying squid than the no-regime model.

*Figure S5:*

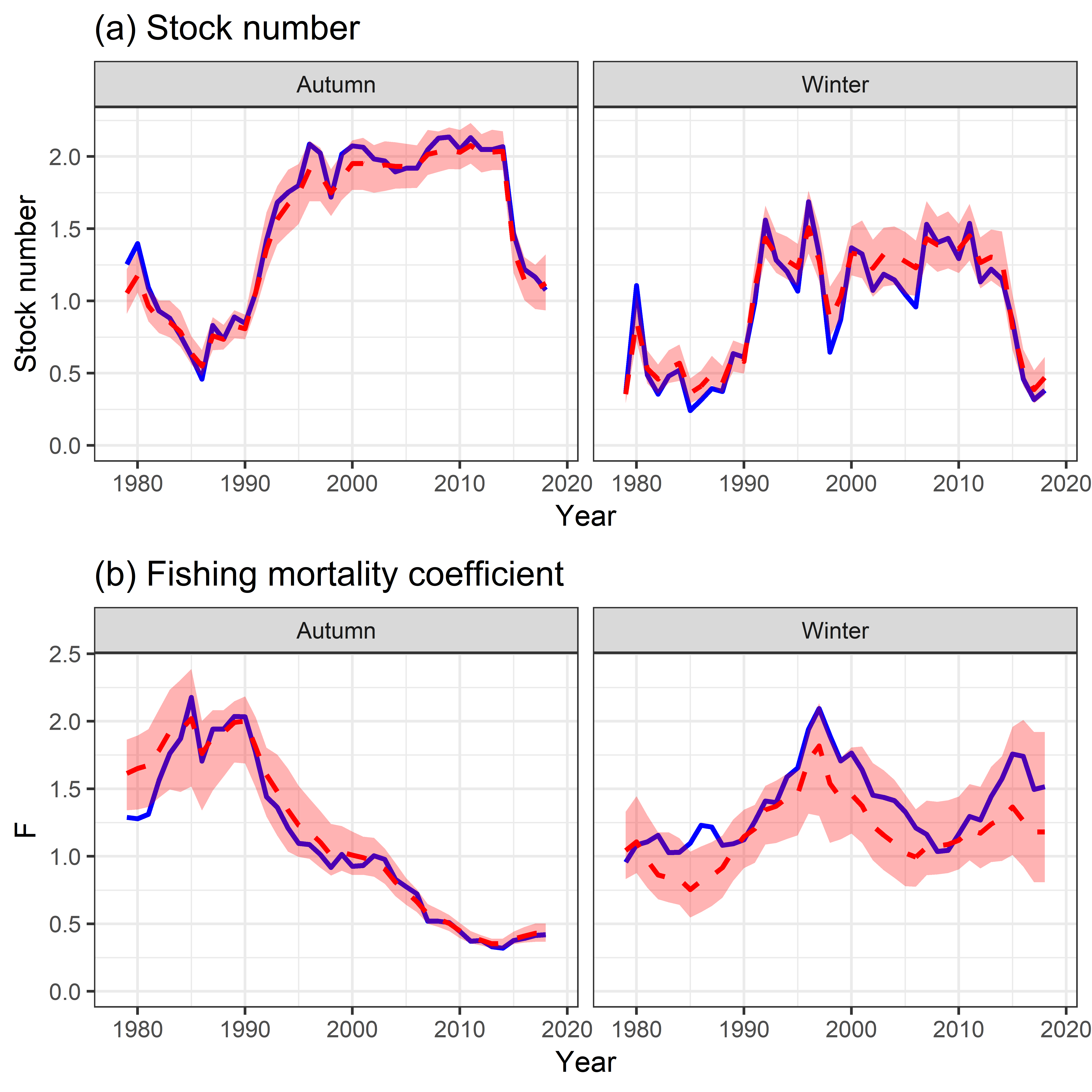

Point estimates (blue) and bootstrapped medians (red, dashed) with 80% confidence interval for (a) stock numbers (billion) and (b) fishing mortality coefficients in the regime-shift model.

*Figure S6:*

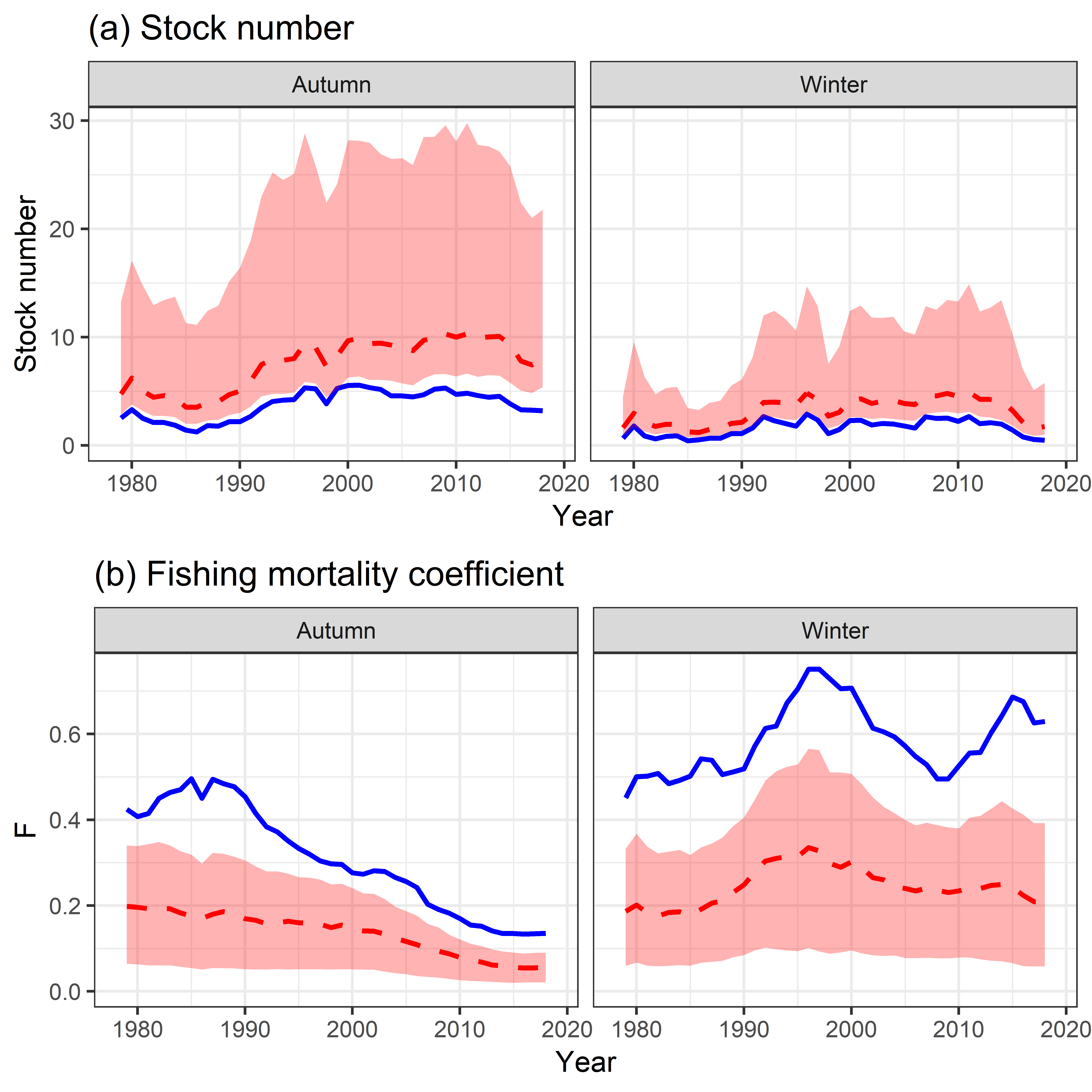

Point estimates (blue) and bootstrapped medians (red, dashed) with 80% confidence interval for (a) stock numbers (billion) and (b) fishing mortality coefficients in the no-regime model.

*Figure S7:*

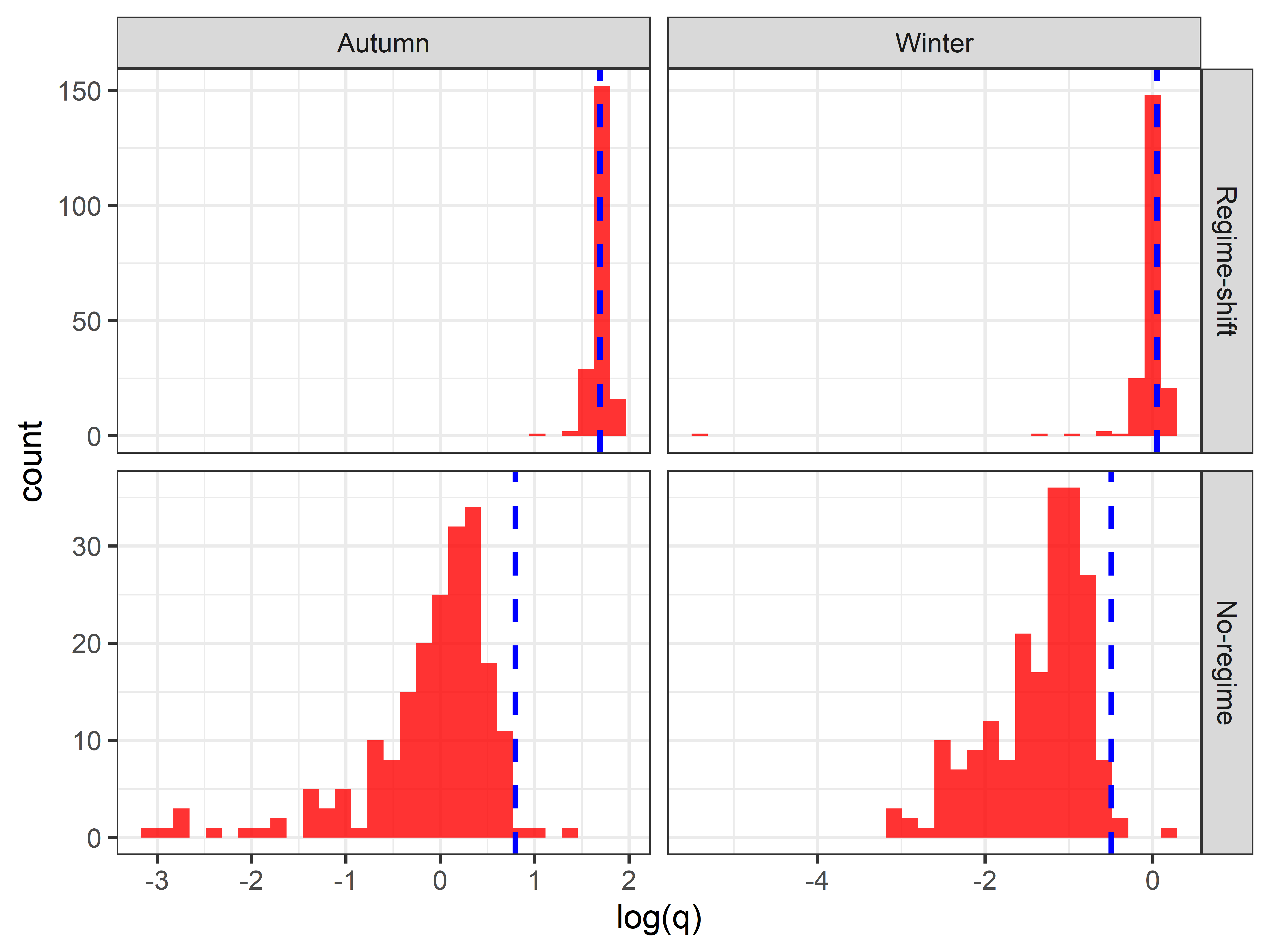

Histogram of log(*q*) in the bootstrap analysis for the autumn-spawning (left) and winter-spawning (right) stocks in the regime-shift (upper) and no-regime (lower) models. The blue dashed lines indicate the point estimates.
